# A simple within-host, between-host model for a vector-transmitted disease

**DOI:** 10.1101/2022.11.28.518273

**Authors:** Mayra Núñez-López, Jocelyn A. Castro-Echeverría, Jorge X. Velasco-Hernández

**Affiliations:** Department of Mathematics, Instituto Tecnológico Autónomo de México, Río Hondo 1, 01080 Ciudad de México, Mexico; Instituto de Matemáticas UNAM Boulevard Juriquilla 3001 Juriquilla 76230 México

**Keywords:** Vector-borne diseases, Multiple time scales, Between-host dynamics, Within-host dynamics, Transmission-clearance trade-off

## Abstract

We present a model that explicitly links the epidemiological Ross-Macdonald model with a simple immunological model through a virus inoculation term that depends on the abundance of infected mosquitoes. We explore the relationship between the reproductive numbers at the population (between-host) and individual level (within-host), in particular the role that viral load and viral clearance rate play in the coupled dynamics. Our model shows that under certain conditions on the strength of the coupling and the immunological response of the host, there can be sustained low viral load infections, with a within-host reproduction number below one that still can trigger epidemic outbreaks provided the between host reproduction number is greater than one. We also describe a particular kind of transmission-clearance trade off for vector-host systems with a simple structure.

## 1. Introduction

Infectious disease dynamics integrates two key processes in the host-parasite interaction. One is the epidemiological process associated with disease transmission, and the other is the immunological process of infection at the individual host level. The transmission of an infectious agent in a population involves various spatial and temporal scales. Specifically, these scales can be broken down into two major groups of phenomena: those that occur at the population scale (epidemic outbreaks) and those that occur within the host (pathogen-immune system interaction). There are a multiplicity of papers of a theoretical nature that have explored this interaction e.g., [1, 2, 3, 4, 5, 6, 7]. The vast majority of these works focus their analysis on directly transmitted diseases based on Kermack-McKendrick type models although there are some studies addressing vector-borne diseases [8, 9]. At the immune system level, the most widely used model, for theoretical purposes, is the one developed for HIV by, for example, [10] that differentiates between target cells, infected target cells, and virions. In particular [5, 6, 7] have looked at the problem of the interplay of between-host, within-host dynamics in an environmentally driven disease framing the equations at the population level as follows

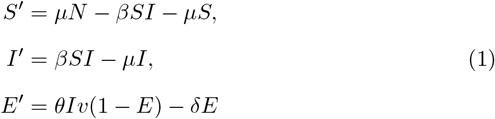

and for the within-host dynamics:

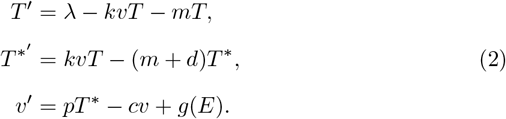

where *S, I*, and *E* denote the susceptible, infectious and polluted environmental compartments with *μ, β*(*v*), *θ* and *δ* being the birth and mortality rate, the infection rate, the shedding rate from infected host deposited in the environment, and *δ* the environmental degradation rate, respectively. For the within-host system, *T, T* ^*^ and *v* represent the target cells, infected target cells and virions respectively; as for the parameters *λ, k, m, d, p* and *c* represent the cell recruitment rate, cell infection rates, cell death rate, virion induced cell death rate, virion production rate and virion clearance rate, respectively. The function *g* represents the inoculum of virions coming from the contaminated environment *E*. In general, it is assumed that 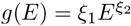 for *ξ*_*i*_ > 1 and *ξ*_2_ ≤ 1 [5, 7].

Feng et al. [7], have generalized results regarding the evolution of virulence by introducing into the host infected class (Eq. 1) a disease-induced death rate. One of these results is that the evolution of virulence in the host population will favor a maximum level of virulence (at the between-host level) if virion production at the cellular level (within-host) is maximal (*p* in Eq. 2) or, alternatively, will favor an intermediate level of virulence if the maximum rate of virion production is large.

In this paper we are interested in exploring between-host, within-host tradeoffs in the context of a vector-borne disease in a vertebrate host. The aim of this article is theoretical. We are interested in exploring the coupled dynamics of within- and between-host dynamics in vector-host transmission systems. As mentioned above, [5, 7] have studied the interaction of these in an infectious disease that has an environmental component represented in Eq. 1. It is through this component that the virus, when interacting with the contaminated environment, is inoculated in the host, thus linking transmission at the population level with infection at the individual level. Here we explore the same type of problem but replace the passive interaction between contaminated environment and host with an insect vector that actively seeks and infects the host. We restrict ourselves only to infection in the mammalian host and do not consider within-vector dynamics.

Vector-borne diseases are a group of diseases of great human importance with nearly half of the world’s population infected with at least one type of vector-borne pathogen [11]. Diseases such as malaria, African trypanosomiasis, Chagas disease and Dengue fever, to mention just a few examples, are serious public health problems in many regions of the world, generating high levels of mortality and morbidity in at-risk populations, which are generally those with the least economic resources and with the least access to adequate public health systems [12]. On the other hand, arthropod-borne diseases are abundant in vertebrates such as horses, cattle and other mammals. Climate change has a direct impact on arthropod vectors (abundance, geographical distribution, and vectorial capacity) [13, 14] producing a reemergence of many infectious diseases both in humans and animals of direct economic importance.

What we seek is to explore the fundamental relationship between reproductive numbers at the population level, at the individual level and, in particular, the role of the within- and between-host system in the epidemic dynamics. As stated above, we postulate a model that explicitly links the epidemiological and immunological dynamics through an inoculation term that depends on the abundance of infected mosquitoes. This approach is based on the idea of separating biological time scales: a fast time scale associated with the within-host dynamics and a slow time scale associated with the epidemiological process. One of the advantages of this approach is that the explicit linkage between the two processes can be established through infected mosquitoes: a bite from an infected mosquito inoculates into the host an extra viral load that connects, within our approach, the population dynamics with the within host dynamics of the disease.

### 2. Model setup

We couple the classical Ross-Macdonald model for a vector-host system coupled with the standard within-host model 2. Hosts are general vertebrate species and the vector is, in general, a mosquito. In the epidemiological model *I* represents the number of infected vertebrate individuals and *Y* represents the number of infected mosquitoes. The variables *T* and *T* ^*^ correspond to the immunological dynamics and represent uninfected and infected target cells, respectively, and *v* represents the virus concentration in plasma of an average infected vertebrate host; *μ* and *δ* represent mortality rates for the vertebrate and mosquito hosts and *γ* is the cure rate of vertebrate hosts. The parameters *α* = *α*(*x*) and *β* = *β*(*x*^*′*^) represent the effective contact rates from mosquito to animal and animal to mosquitoes and are assumed to depend, in general, on some measure of infectiveness either in the mosquito or vertebrate host, respectively. The most common assumption for these functions [15, 16] is that for *x, x*^*′*^ ∈ [0, ∞) they satisfy *α*(*x*), *β*(*x*^*′*^) ≥ 0, *α*^*′*^ (*x*), *β*^*′*^ (*x*^*′*^) *>* 0, and *α*^*′ ′*^ (*x*), *β*^*′ ′*^ (*x*^*′*^) ≤ 0. In our model (Eq. 3 below), *β*(*x*^*′*^) is the biting rate that transmit the disease from an infected host with infectiousness *x*^*′*^ to a susceptible mosquito. A hypothesis of our model is that the biting rate from the infected mosquito to a susceptible host, *α*(*x*) will be proportional to *β*(*x*^*′*^). Some evidence supporting this hypothesis is in the work of Tesla et al. [17] who report that, for Zika, increasing viral dose in the blood-meal significantly increases the probability of mosquitoes becoming infected and becoming infectious. This hypothesis simplifies our model because then we do not have to follow the fate of the viral load in the mosquito. In summary, the rationale of this assumption is that a high viral load in the vertebrate host will generate a high viral load infection in the mosquito that, in turn, will produce a high effective biting rate of infected mosquitoes to the vertebrate host. For the within-host system, *λ* represents the recruitment rate of healthy target cells, *m* the natural mortality rate of target cells, *k* the cell-infection rate, *d* the virus-induced cell death, *p* the virus proliferation rate per infected cell, *c* the viral clearance rate and *g* = *g*(*y*) is an inoculation term that depends on the abundance of infected mosquitoes *y*. Let *α*(*x*) = *ab*(*x*) where *a* is the biting rate and *b*(*x*) is the probability of vertebrate infection per bite; likewise, *β*(*x*^*′*^) = *aϕ*(*x*^*′*^) where *ϕ*(*x*^*′*^) is the probability of mosquito infection per bite. The equations for the between-host system are a variant of the so-called Ross-Macdonald equations:

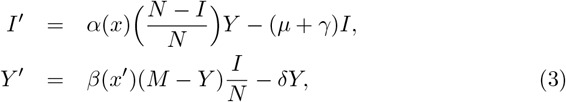

The equations for the within-host dynamics are now:

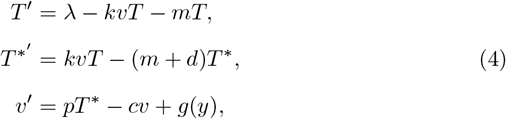

where *N* and *M* stand for the total constant populations of vertebrate host and mosquito, respectively. Normalizing Eq. (3) by defining *i* = *I/N* and *y* = *Y/M*, and defining *q* = *M/N* we can rewrite them as

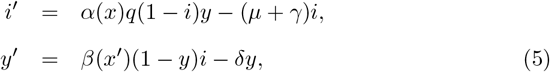

This is the epidemiological model that will be studied below. In order to link the abundance of infected mosquitoes with the infection process at the individual level, we assume that infected mosquitoes directly correlate with within-host level of infected target cells. This biological consideration suggest that the function *g* should have the following properties: *g*(*y*) ≥ 0, *g*(0) = 0, *g*^*′*^ (*y*) *>* 0 and *g*^*′ ′*^ (*y*) ≤ 0.

In general, as in [5], we must take *g*(*y*) = *ry*^*s*^ with *r, s >* 0. In the next section we restrict our analysis to the case *s* = 1 as our aim is to illustrate the framework of linking within- and between-host dynamics for viral load-dependent contact rates. The inclusion of the inoculation rate *g*(*y*) is key for linking the within-host dynamics to the between-host dynamics and replaces the environmental inoculum described in [5, 6, 7].

### 3. Model analysis

An important biological feature of this coupled system is that the within-host dynamics occurs on a faster time scale than the dynamics of the between-host and the environment. This multiple time-scale allows us to study the mathematical properties of the model by analyzing the fast- and slow-systems determined by the two time scales. As evidence that supports this analysis we can cite [18] who reports on the duration of DEN-1 viremia in a clinical study. According to this author, the duration of viremia ranged from 1 to 7 days (mean, 4.5 days; median, 5 days) with viremias of primary infection lasted more compared to secondary infections: the mean duration of viremia for all patients experiencing a primary dengue virus infection was of 5.1 days versus 4.4 days for those with a secondary dengue virus infection. In contrast Dengue outbreaks last several months or, in endemic situations, transmission takes place over the years as reported in [19] or the statistics provided by PAHO, among many other sources. We would like to decouple model (3, 4) with respect to time. As done in [5, 6, 7] we would like to separate slow and fast subsystems corresponding to either of the between-host (epidemiological) or the within-host (immunological) models.

#### 3.1. Summary of results for the fast subsystem

The fast system has been analyzed by [5, 7] for the case of an environmentally-driven infectious disease. Their results have immediate applicability to our case. In this subsection we briefly summarize them. The within-host dynamics (4) can be considered the fast system where the variable *y* can be treated as a constant (i.e. it is not changing with time on the fast time scale). In our case (Eq. 4) when *g*(*y*) = 0, the system always has the infection-free equilibrium) 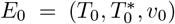 where 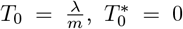. Let *R*_*v*_(*y*) denote the within-host reproduction number, which is a function of the density of infected mosquitoes, and define *R*_0*v*_ = *R*_*v*_(0) given by

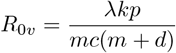

as the basic reproduction number of the uncoupled fast (within-host) system. As in Feng et al. [6], *R*_*v*_(*y*) when *y >* 0 is given by

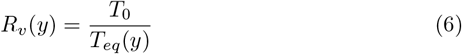

where we take the biological feasible solution to be (cf Feng et al. [6]):

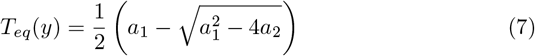

with

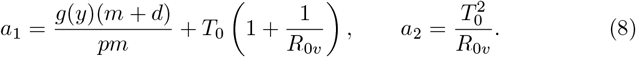

*R*_0*v*_, the within-host reproduction number does not depend on *y* but the reproductive function *R*_*v*_(*y*) depends on the magnitude of *R*_0*v*_. Such a dependence is illustrated in Fig 1. This figure plots the curves *R*_*v*_(*y*) for different *R*_0*v*_ values, where we have used a linear function for *g*(*y*) = *ry* with *r* constant. Following Feng et al. [6], we know that given *R*_0*v*_ *>* 1, lim_*y*→0_ *R*_*v*_(*y*) = *R*_0*v*_; otherwise if *R*_0*v*_ < 1, then lim_*y*→0_ *R*_*v*_(*y*) = 1 (see the upper right corner of Fig. 1). *E*_0_ is locally asymptotically stable if *R*_0*v*_ < 1 and unstable if *R*_0*v*_ *>* 1.

**Figure 1:**
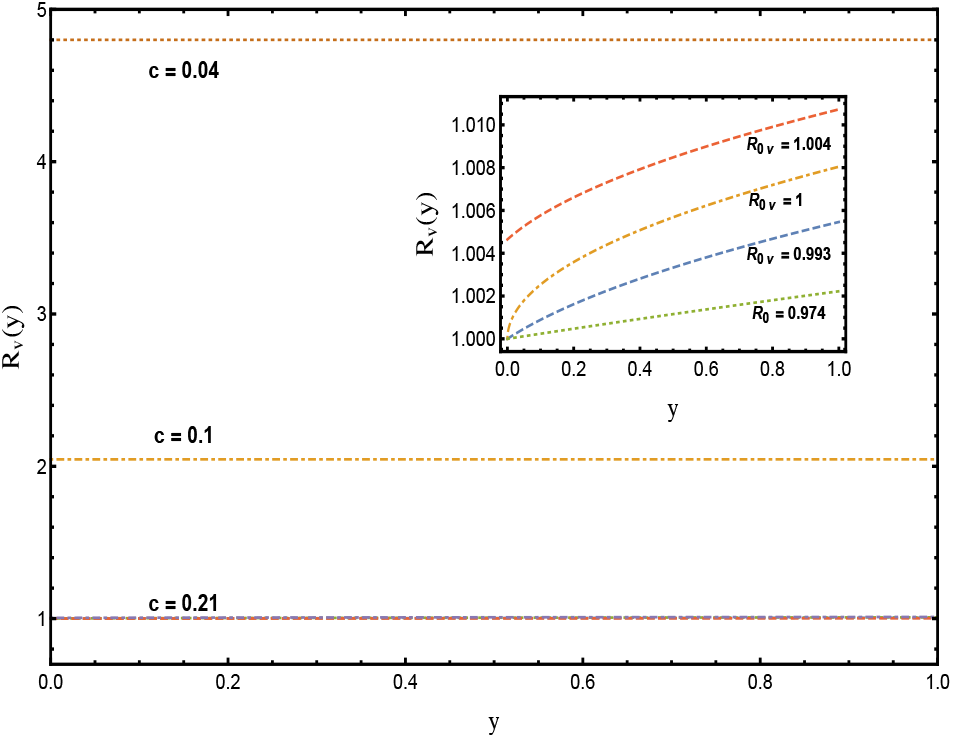
Within-host reproduction number *R*_*v*_ (*y*) as functions of vector prevalence *y*. Parameters *λ* = 5000, *m* = 0.311, *d* = 0.01, *p* = 10^4^, *r* = 10000, *k* = 4.08410^−10^.

In Appendix A, we present the global stability of the disease-free equilibrium point *E*_0_ with *g*(*y*) = 0 for the within-host dynamics.

There exists a unique endemic equilibrium *E*_*f*_ = (*i*^*^, *y*^*^) with *i*^*^ *>* 0, *y*^*^ *>* 0 of the fast system if and only if *R*_0*v*_ *>* 1. Following [5] the endemic equilibrium *E*_*f*_ is locally asymptotically stable whenever *R*_0*v*_ *>* 1.

#### 3.2. The slow subsystem

Let *x*^*′*^ = *T* ^*^(*t*)*/T*_0_, the proportion of infected target cells at time *t*, a measure of the infectiousness of the vertebrate host. Let

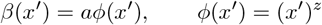

with 0 *<z* and *a >* 0, the biting rate. In the case of vertebrate infections, these depend on the infectiousness of the mosquito bite. Since we are not following the within-mosquito dynamics, we will let *b*, the probability of infection form mosquito to vertebrate host to be a free parameter. The basic reproduction number is then

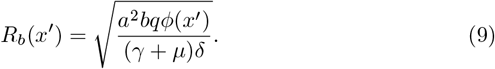

Note that if *x*^*′*^ = 1 then

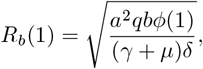

is the maximum biologically feasible reproduction number as a function of host infectiousness *x*^*′*^. When *R*_*b*_(*x*^*′*^) *>* 1, the (between-host) endemic equilibrium point can exists and be found explicitly:

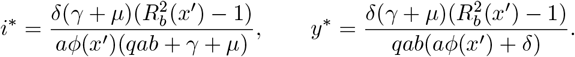

Both of these coordinates depend on *x*^*′*^ and will render the between-host endemic equilibrium only when 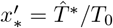, the equilibrium infected target cell infection. An alternative way of looking at the between-host endemic equilibrium is the following. The endemic equilibrium point (slow subsystem) (*i*^*^, *y*^*^) is located on the intersection of the zero isoclines of the between-host equations (for constant within-host dynamics). Explicitly, these are

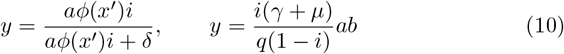

The intersection exists with positive *i* whenever *R*_*b*_(*x*^*′*^) *>* 1 which is the standard condition for the existence of an endemic equilibrium point in the Ross-Macdonald model. However, in this case, our equilibrium will be located on the line that describes this intersection as function of the parameter *x*^*′*^ (see Figure 2) and it will be determined when 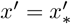 implying that *R*_*b*_(*x*^*′*\*^) = *R*_0*b*_ ≤ *R*_*b*_(1), i.e., the between-host basic reproduction number is bounded by the maximum of the between-host reproduction function. We now proceed to characterize the within-host endemic equilibrium, particularly how its state variables depend on the (population level) mosquito abundance. In this, we follow Feng et al. [5]. The epidemiological and within-host subsystems are linked through the abundance of infected mosquitoes in terms of *R*_*v*_(*y*) (Eq. 6). Assume *R*_*v*_(*y*) *>* 1 and that the fast system is at its stable nontrivial equilibrium 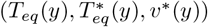 given by (6, 7 and 8), where 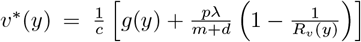 is indicated in Appendix B.

**Figure 2:**
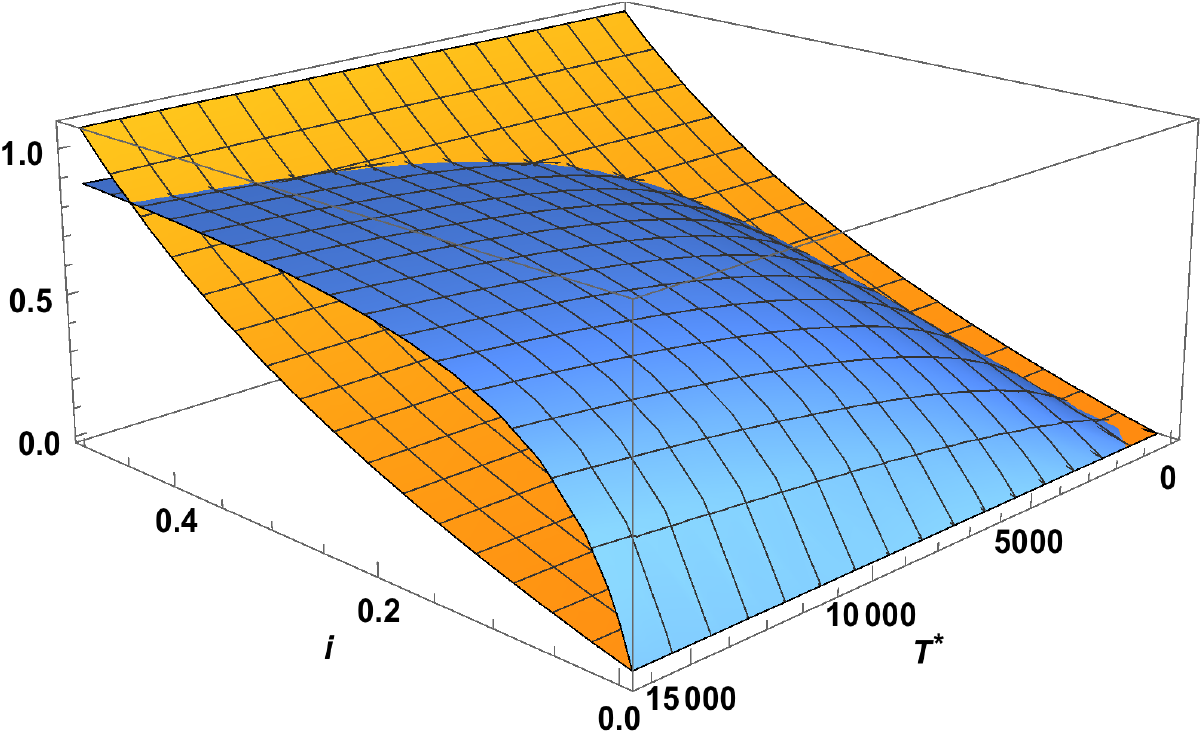
3D representation of the zero-isoclines for the between-host model as a function of *x*^*′*^, the proportion of infected target cells. *x*-axis is *i, y* axis is *T* ^*^ with *z* = 0.8. The blue shaded region describes the dynamic transcritical bifurcation that appears after *T* ^*^ reaches the critical value such that *R*_*b*_(*T* ^*^) = 1.

Note that

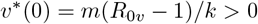

when *R*_0*v*_ *>* 1. The viral load at equilibrium depends now on *y* and to have *v*^*^(*y*) *>* 0, it is required that the within-host reproduction function *R*_*v*_(*y*) = 2*Q >* 1 where

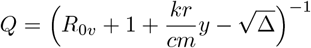

and

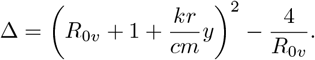

Biological feasibility dictates that Δ, *Q >* 0. This is satisfied if *R*_0*v*_ *>* 1 since *R*_*v*_(*y*) is an increasing function of *y*.

### 4. Linking time scales

The Jacobian of the whole coupled system is

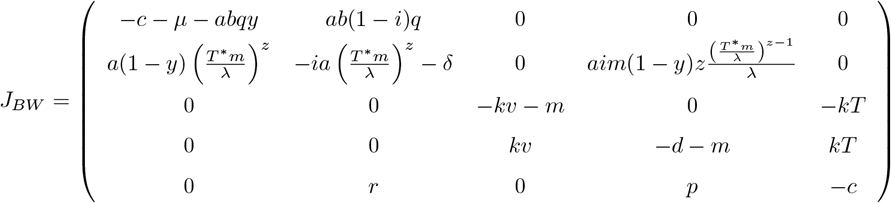

At the disease-free equilibrium *E*_0_ = (0, 0, *λ/m*, 0, *v*^*^(0)) the Jacobian is

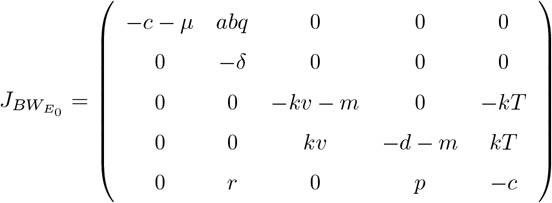

Note from 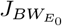 that the components of the between-host reproduction number, namely the biting rates, play a role on the stability of *E*_0_ through the proportion of infected target cells *T* ^*^*m/λ* once the within-host infections starts to grow when *R*_0*v*_ *>* 1. Also, since this condition implies that lim_*y*→0_ *v*(*y*) *>* 0 then necessarily lim_*y*→0_ *T* ^*^(*y*) *>* 0 too.

### 5. Conditions for a disease outbreak

#### 5.1. The epidemic system

The existence of an epidemic outbreak depends on the strength of the infection at the within-host level measured by the within-host reproduction number when *R*_0*v*_ *>* 1. The between-host reproduction number *R*_*b*_(*x*^*′*^) will be greater than one only until enough infection has accumulated so as to sufficiently increase the ratio *x*^*′*^(*t*) = *T* ^*^(*t*)*m/λ*. When 0 *<R*_*b*_(*x*^*′*^) < 1 the only between-host equilibrium point that exists is the disease-free equilibrium which is asymptotically stable. *R*_*b*_(*x*^*′*^) *>* 1 requires the average individual in the population to have an active (within-host) viral infection but the transmission efficacy will not be large enough so as to trigger an epidemic until *R*_*b*_(*x*^*′*^) = *R*_0*b*_. We can give a more detailed description of the dependence of the between-host equilibrium state and the within-host dynamics. First, there exists a critical value of 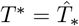 where 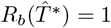. In Figure 2 we plot the intersection of Eq. (10) to show how the existence of an endemic equilibrium depends on *T* ^*^ and *i*. As *T* ^*^ increases above 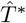, the boundary between the two colored regions shown in the figure is the line that contains the feasible endemic equilibrium that is realized when *T* ^*^ reaches its steady-state. A second important feature is associated with the contact rate *β*(*x*^*′*^) = *aϕ*(*x*^*′*^), where *ϕ*(*x*^*′*^) = (*x*^*′*^)^*z*^. Figure 3 shows the intersection of the two isoclines Eq (10) that give the feasible endemic equilibrium but as functions of *x*^*′*^ and *z*, the exponent of the probability of infection *ϕ*(*x*^*′*^). Large values of *z* prevent the existence of an endemic between-host equilibrium point, whereas for *z* ≤ 1 the endemic equilibrium always exist. So concave probabilities of infection always generate an endemic state provided *R*_*b*0_ *>* 1 while convex ones do not. A third observation is that we can expect a time-delay of variable duration occurring between the crossing of the threshold *R*_*b*_(*T* ^*^) = 1 and the time when the epidemic outbreak will occur and will send the between-host system to its endemic state, i.e., when *R*_*b*_(*T* ^*^) = *R*_0_. This delay appears because of the dynamic nature of our contact rate parameters that depend on the within-host dynamics.

**Figure 3:**
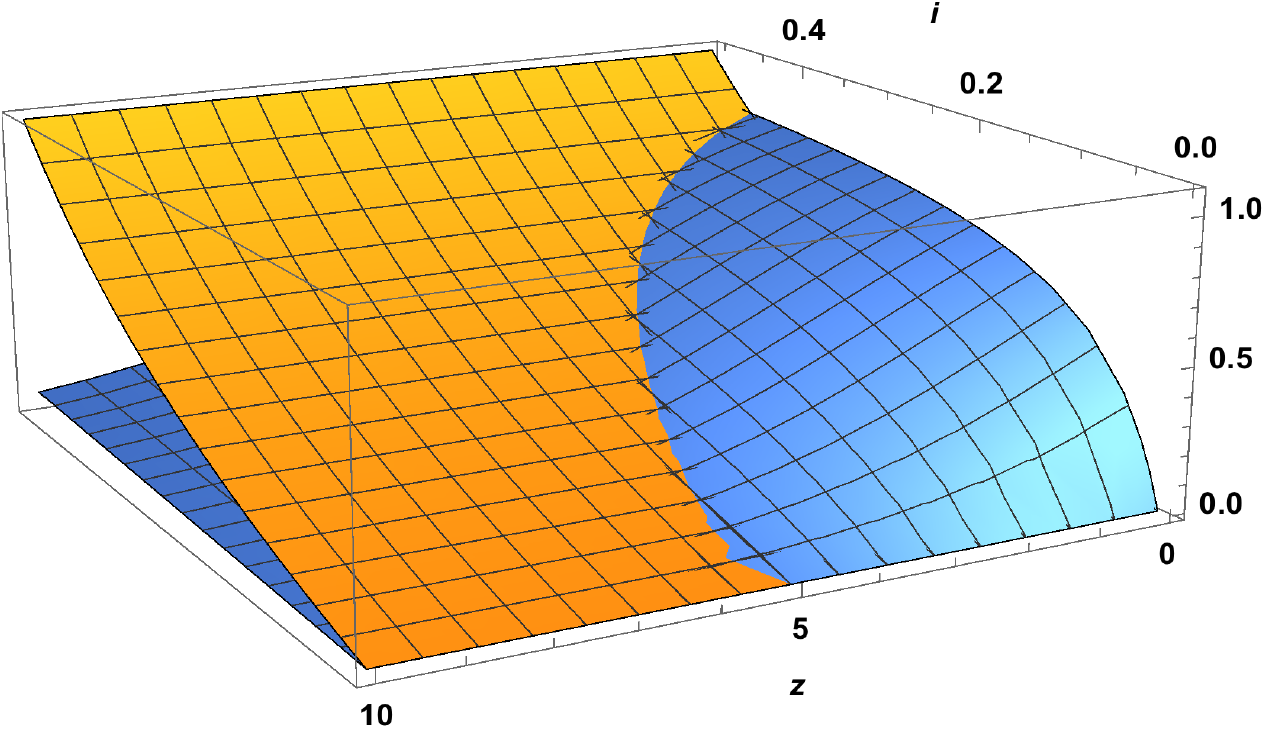
3D representation of the zero-isoclines for the between-host model as a function of *x*^*′*^, the proportion of infected target cells. *x*-axis is *i, y* axis is *z*. The blue shaded region describes the transcritical bifurcation that appears after *T* ^*^ reaches the critical value such that *R*_*b*_ (*T* ^*^) = 1.

#### 5.2. The full coupled system

We look now at the role of virulence, measured by our variable *x*^*′*^, on the dynamics of our system. First, we make the reasonable assumption that the recovery rate *γ* is related to the viral clearance rate *c* in a very specific way. We postulate that the recovery rate is not constant but satisfies

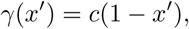

implying that large virulence is associated with chronic disease with practically no recovery, and low virulence makes the recovery rate *γ* approximately equal to the clearance rate *c*. Recall that *R*_*b*_(*x*^*′*^) is given by Eq. (9). The endemic equilibrium for the host population, on the other hand, has the formula

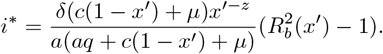

We can easily prove that, as a function of *x*^*′*^, *i*^*^(*x*^*′*^) is a monotonically increasing function, and that *R*_*b*_(*x*^*′*^) is concave, if *z <* 1 and convex is *z >* 1. Also *i*^*^ is biologically feasible only for 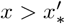 where 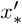 is the proportion of infected target cells that results in 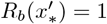. Moreover, for the same value of *x*^*′*^, transmission probabilities with *z >* 1 produce lower levels of endemicity than for *z <* 1. The temporal dynamics of the coupled system is depicted in (Table 1 top to bottom). In all these simulations, we are assuming that the recovery rate of infected individuals is of the form *γ*(*x*^*′*^) = *c*(1 − *x*^*′*^) with *x*^*′*^ = *T* ^*^*/T*_0_ and, also, we have set the baseline clearance rate to *c* = 0.14 or, equivalently, a duration of viremia lasting 7 days. Seven days is then, the shortest recovery time. Since we are using the same within-host parameters in all runs, the behavior of *x*^*′*^ in all cases is the same as can be seen in Table 1b. The observed delay in the onset of the epidemic (Table 1a) at the between-host level is associated with the particular shape of the probability of infection *ϕ*(*x*^*′*^) = (*x*^*′*^)^*z*^ (see Eq.9) from mosquito to host and the ratio of mosquito numbers to host numbers. The rows of Table 1 correspond to different values of the parameter *z*. Top and middle rows correspond to *z <* 1 and the bottom row to *z* = 1. We can see that for the same within-host dynamics, slowly growing transmission probabilities (Table 1 top row) provide a earlier outbreak than faster growing ones (Table 1 middle row). However, this effect can be modified by the magnitude of the product *bq* = *bM/N* the effective ratio of mosquito to host (Table 1 bottom row). Table 1c shows the relative magnitud of the within-host (constant) reproduction number and the between-host reproduction function as *x*^*′*^ changes. Finally, Table 1d shows that when *R*_*b*_(*x*^*′*^) *<R*_0*v*_ the mosquito infection at the population level, is slower than the infection of target cells (middle and bottom rows). If *R*_*b*_(*x*^*′*^) *> R*_0*v*_ the above condition still holds but both time scales are then very similar.

**Table 1:**
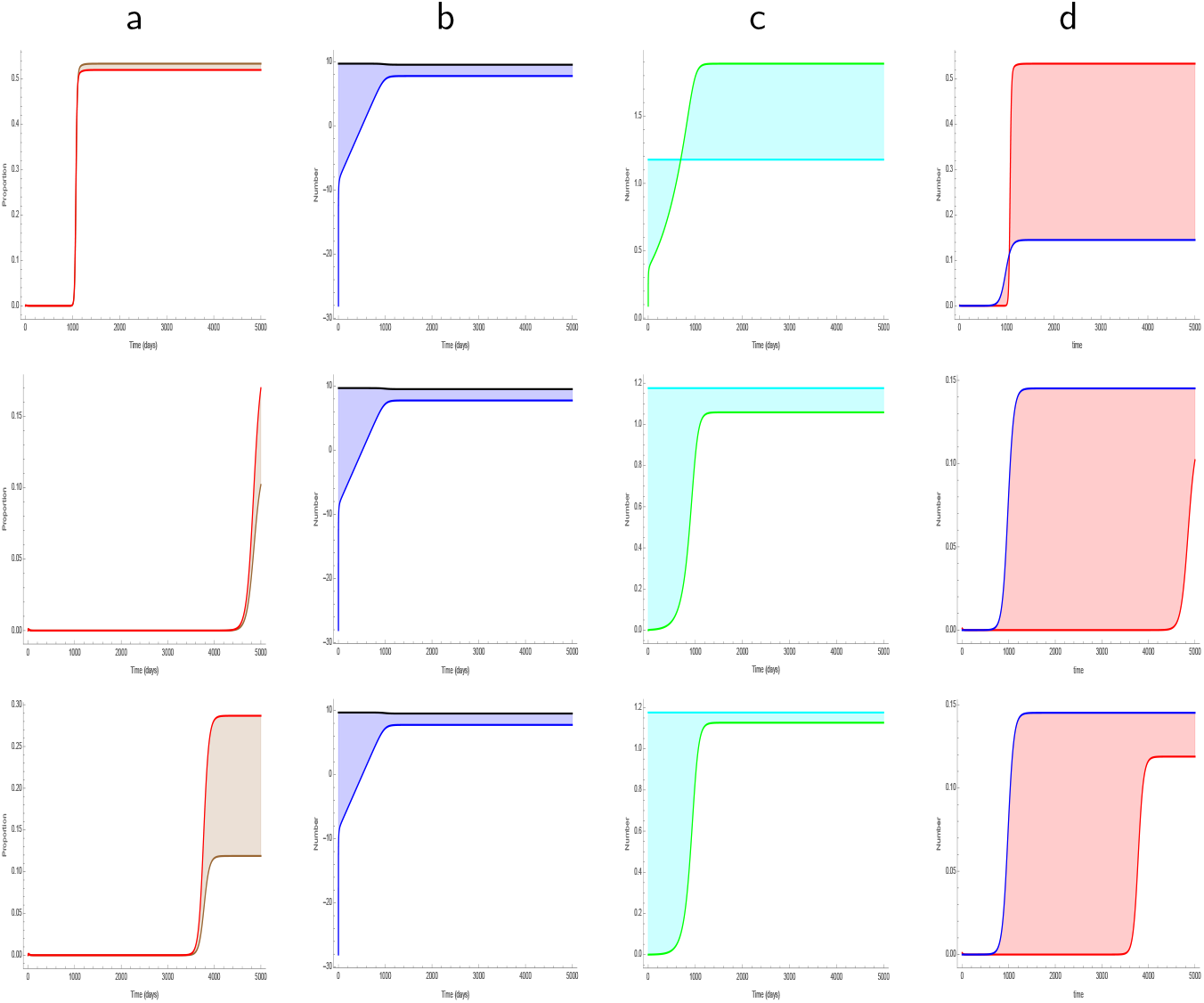
Dynamic behaviour of the full within-host, between-host model. Parameters for the within-host system are as in Table 2. For the between-host system, *a* = 0.162, *γ* = *c* = 0.14 and *δ* = 0.05. Rows correspond to different values of *z*. Top, *z* = 0.2, middle *z* = 0.8 both with *q* = 1.5 mosquitos per host; and bottom *z* = 1 with *q* = 2.5 mosquitoes per host. Columns show a) the prevalence of infected mosquitoes (brown) and vertebrate hosts (red); b) in logarithmic scale the density of naive (black) and infected (blue) target cells, c) the within-host reproduction number *R*_0*v*_ (cyan), the between-host reproduction function *R*_*b*_ (*x*′) (green); the proportion of infected target cells *x* (blue) and the proportion of infected mosquitoes *i* (red).

Finally, looking closer to the clearance rate *c* we can say that, in general 0 *<c*_*_ ≤ *c* ≤ *c*^*^ where *c*^*^ is the value of *c* for which *R*_0*v*_ = 1. As *c* → 0 the within-host reproduction number *R*_0*v*_ tends to infinity but *R*_*v*_(*y*) ceases to be a real number and the ODE system breaks down. In summary, the upper bound is determined solely by the within-host dynamics *R*_0*v*_, but a very large residence time 1*/c* is biologically unfeasible given the mathematical model we are proposing. Table 2 shows how *R*_0*v*_, *v*^*^(*y*) and *R*_*v*_(*y*) depend on the parameter *c*. We have arbitrarily selected three regions in this curves. It is clear that large or intermediate values of *c* (as described in the figure caption) are biologically feasible giving a reasonable magnitude range for the within-host reproduction number (e.g., *R*_0*v*_ < 4). A large *c* describes a short viremia period while a small *c* describes a long one. Our results thus indicate that, for the kind of interaction described by our model, short or intermediate viremia duration are biologically more feasible than long ones. On the other hand, very short viremias render *R*_0*v*_ < 1 and the whole coupled system breaks down (mathematically, solution no longer exist).

**Table 2:**
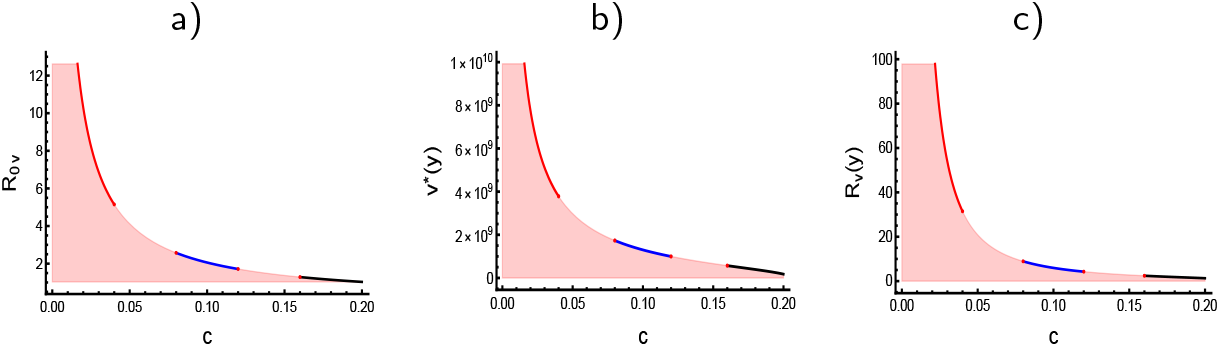
Key within-host functions and their dependence on *c*, the clearance rate. Parameters are *k* = 4.08410^−10^, *λ* = 5000, *m* = 0.311, *d* = 0.01, *p* = 10^4^, *r* = 10000. For these particular parameters, *c*^*^ ≈ 0.205847. The colored segments represent examples of clearance rates of short duration (black), medium duration (blue) and long duration (red). To the left of the blue region, the values of all of these functions are biologically unfeasible. (a) Within-host reproduction number *R*_0*v*_; (b) Approximate viral load function *v*^*^(*y*); (c) Reproduction function *R*_*v*_ (*y*), Eq.(6)

The within-host and between-host population processes are closely coupled as can be seen in the timing where equilibria for both subsystems is reached (Table 1 columns a, b and d). However, the between-host reproduction function approaches its limit *R*_*b*_(*T* ^*^) at different speeds depending on the magnitude of *z*. The epidemic outbreak will be triggered when the within-host system reaches its equilibrium state regardless of how large is the within-host infection while approaching it. So, our model indicates that transmission at the population level is feasible but cannot be realized until the average infection conditions of individuals reach their corresponding equilibrium. Therefore, the reproduction number of the between-host system is an indicator of an epidemic outbreak that will be occurring later in time, depending upon the magnitude of *T* ^*^. This is one of the explicit links of the population level reproduction number and the dynamics of the within-host infection.

## 6. Conclusion

The dynamics of infections diseases is driven by two processes: the epidemiological process occurring at the population level and the immunological process within the host. Many existing models in the context of a vector-borne disease, approach these two process as decoupled systems. In this paper we have linked them using a simple model based on two classical well-known equations: the Ross-Macdonald model and a basic virus-cell interaction model. We demonstrate our framework by using as a simplified model system for a general vector-borne disease in a vertebrate host. Naturally, in these diseases the vector plays a major and determinant role in transmission. This model produces a clearance-transmission trade-off where viral load is an increasing function of viremia duration. This results is contrary to the results on Dengue reported by [20], where short viremias have larger viral loads than long viremias. Due to the way the within-host dynamics is modelled and the resulting form that the within-host reproduction number takes, large *c* (short viremias) reduce the magnitude of the reproductive number and therefore, generate lower viremias than when *c* is small (long viremias). For Dengue disease, [20] use a more detailed model carefully adapted to Dengue viral dynamics that is able to capture dynamical characteristics that our simple model cannot achieve. Our simple model does not consider any specific mechanisms of activation of the innate and adaptive immune responses and thus our results cannot directly be compared to those in [20]. However, results on malaria [21] may seem to agree with the relation of clearance and pathogen load that our model produces. In this work it is clear that the length of pathogen clearance time is positively associated with higher concentrations of parasites. For Zika, [22] reports relatively long viremias in whole blood samples in human hosts, of more than 26 days, while in macaques [23], the highest viremia was reported for intermediate duration (in macaques the viremia length ranges form 2 to 7 days); for Chikungunya, [24], the higher frequency of high viremia in human hosts occurred also on the 7 day of symptom onset (symptom onset occurs in the interval 1-20 in this study). The model we develop and analyze in this work integrates in a simple and direct manner, the interplay of epidemiological dynamics and within-host immune-virus interaction dynamics. The model focuses in a general, classical approach to approximating the dynamics of vector-borne diseases and immune system dynamics on vertebrate hosts. The conclusions therefore, are also of a general nature and only describe broad patterns of interaction.

## Acknowledgemets

MNL acknowledges the financial support from the Asociación Mexicana de Cultura, A.C.; JACE and JXVH acknowledge support from grants UNAM PA-PIIT IN115720 and IV100220. JXVH developed part of this work during his sabbatical year at Departmento de Matemáticas ITAM and the Simons Institute, University of Berkeley. For his stay at the latter, support from PASPA-UNAM fellowship is acknowledged.

## Appendix A

*Global stability of the disease-free equilibrium point* 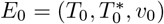 *for the within-host dynamics (fast subsystem) given by system (4) with g*(*y*) = 0.

Let 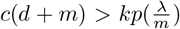, i.e. *R*_0*v*_ < 1 and 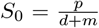, then the critical point *E*_0_ is globally asymptotically stable.

In order to prove the asymptotic stability of *E*_0_, consider the Lyapunov function

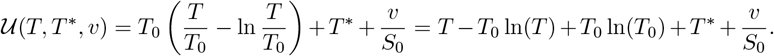

Then from the last expression

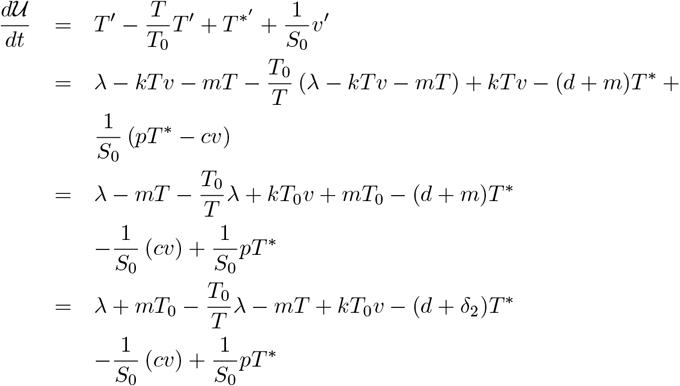

Substituting *mT*_0_ = *λ* in the second term, *m* = *λ/T*_0_ in the forth term and *S*_0_ = *p/*(*d* + *m*) in the last one we get

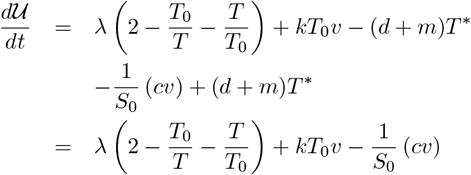

A further simplification yields

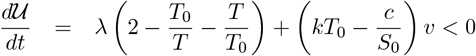

The last inequality follows from the hypothesis and the inequality of the geometric and arithmetic means.

## Appendix B

Here we give the full expression of the within-host reproduction function *R*_*v*_(*y*) that appears in expression *v*^*^(*y*)

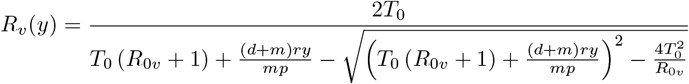

## Author’s contributions

JXVH conceived the project; MNL, JACE and JXVH performed the analyses. MNL and JXVH wrote the manuscript. All authors discussed and revised the manuscript.

## Bibliography

[1] M. A. Gilchrist, A. Sasaki, Modeling host−parasite coevolu-tion: A nested approach based on mechanistic models, Journal of Theoretical Biology 218 (3) (2002) 289–308. doi:https://doi.org/10.1006/jtbi.2002.3076. URL https://www.sciencedirect.com/science/article/pii/S0022519302930766

[2] M. A. Gilchrist, D. Coombs, Evolution of virulence: Interde-pendence, constraints, and selection using nested models, Theo-retical Population Biology 69 (2) (2006) 145–153. doi:https://doi.org/10.1016/j.tpb.2005.07.002. URL https://www.sciencedirect.com/science/article/pii/S0040580905000961

[3] N. Mideo, S. Alizon, T. Day, Linking within- and between-host dynamics in the evolutionary epidemiology of infectiousdiseases, Trends in Ecology Evolution 23 (9) (2008) 511–517. doi:https://doi.org/10.1016/j.tree.2008.05.009. URL https://www.sciencedirect.com/science/article/pii/S0169534708002188

[4] A. E. S. Almocera, V. K. Nguyen, E. A. Hernandez-Vargas, Multiscalemodel within-host and between-host for viral infectious diseases, Jour-nal of Mathematical Biology 77 (4) (2018) 1035–1057. doi:10.1007/s00285-018-1241-y. URL https://doi.org/10.1007/s00285-018-1241-y

[5] Z. Feng, J. Velasco-Hernandez, B. Tapia-Santos, M. C. A. Leite, A model for coupling within-host and between-host dynamics in an infectious dis-ease, Nonlinear Dynamics 68 (3) (2012) 401–411.

[6] Z. Feng, J. Velasco-Hernández, B. Tapia-Santos, A mathematical model forcoupling within-host and between-host dynamics in an environmentally-driven infectious disease, Mathematical Biosciences 241 (1) (2013) 49–55. doi:https://doi.org/10.1016/j.mbs.2012.09.004. URL https://www.sciencedirect.com/science/article/pii/S0025556412001836

[7] Z. Feng, X. Cen, Y. Zhao, J. X. Velasco-Hernandez, Cou-pled within-host and between-host dynamics and evolution ofvirulence, Mathematical Biosciences 270 (2015) 204–212, from Within-Host Dynamics to the Epidemiology of Infectious Disease. doi:https://doi.org/10.1016/j.mbs.2015.02.012. URL https://www.sciencedirect.com/science/article/pii/S0025556415000553

[8] H. Gulbudak, V. L. Cannataro, N. Tuncer, M. Martcheva, Vector-BornePathogen and Host Evolution in a Structured Immuno-EpidemiologicalSystem, Bulletin of Mathematical Biology 79 (2) (2017) 325–355. doi:10.1007/s11538-016-0239-0. URL https://doi.org/10.1007/s11538-016-0239-0

[9] M. Martcheva, N. Tuncer, Y. Kim, On the principle of host evolutionin host−pathogen interactions, Journal of Biological Dynamics 11 (sup1) (2017) 102–119, pMID: 26998890. arXiv:https://doi.org/10.1080/17513758.2016.1161089, doi:10.1080/17513758.2016.1161089. URL https://doi.org/10.1080/17513758.2016.1161089

[10] R. M. Ribeiro, A. S. Perelson, The analysis of hiv dynamics using math-ematical models, AIDS and Other Manifestations of HIV Infection 905 (2004) 912.

[11] I. of Medicine, Vector-Borne Diseases: Understanding the Environmental, Human Health, and Ecological Connections: Workshop Summary, The National Academies Press, Washington, DC, 2008. doi:10.17226/11950.

[12] S. Bhatt, P. W. Gething, O. J. Brady, J. P. Messina, A. W. Farlow, C. L. Moyes, J. M. Drake, J. S. Brownstein, A. G. Hoen, O. Sankoh, M. F. Myers, D. B. George, T. Jaenisch, G. R. W. Wint, C. P. Simmons, T. W. Scott, J. J. Farrar, S. I. Hay, The global distribution and burden of dengue, Nature 496 (7446) (2013) 504–507. doi:10.1038/nature12060.

[13] F. Beugnet, M. Jean-Lou, Emerging arthropod-borne diseases of companion animals in Europe, Veterinary Parasitology 163 (4) (2009) 298–305. doi:10.1016/j.vetpar.2009.03.02.

[14] C. A. Hill, F. C. Kafatos, S. K. Stansfield, F. H. Collins, Arthropod-borne diseases: vector control in the genomics era, Nature Reviews Microbiology 3 (3) (2005) 262–268. doi:10.1038/nrmicro1101.

[15] H. J. Bremermann, H. R. Thieme, A competitive exclusion principle for pathogen virulence, ournal of Mathematical Biology 27 (1989) 179–190. doi:10.1007/BF00276102.

[16] S. Lion, J. A. J. Metz, Beyond r0 maximisation: On pathogen evolution and environmental dimensions, Trends in Ecology and Evolution 33 (2018) 458–473. doi:10.1016/j.tree.2018.02.004.

[17] B. Tesla, L. R. Demakovsky, H. S. Packiam, E. A. Mordecai, A. D. Rodríguez, M. H. Bonds, M. A. Brindley, C. C. Murdock, Estimating the effects of variation in viremia on mosquito susceptibility, infectious-ness, and r0 of zika in aedes aegypti, PLoS Neglected Tropical Diseases 12. doi:10.1371/journal.pntd.0006733.

[18] D. W. Vaughn, S. Green, S. Kalayanarooj, B. L. Innis, S. Nimmannitya, S. Suntayakorn, T. P. Endy, B. Raengsakulrach, A. L. Rothman, F. A. Ennis, A. Nisalak, Dengue Viremia Titer, Antibody Response Pattern, and Virus Serotype Correlate with Disease Severity, The Journal of Infectious Diseases 181 (1) (2000) 2–9. arXiv:https://academic.oup.com/jid/article-pdf/181/1/2/17994503/181-1-2.pdf, doi:10.1086/315215. URL https://doi.org/10.1086/315215

[19] N. Haider, Y.-M. Chang, M. Rahman, A. Zumla, R. A. Kock, Dengueoutbreaks in bangladesh: Historic epidemic patterns suggest earliermosquito control intervention in the transmission season could re-duce the monthly growth factor and extent of epidemics, Current Research in Parasitology Vector-Borne Diseases 1 (2021) 100063. doi:https://doi.org/10.1016/j.crpvbd.2021.100063. URL https://www.sciencedirect.com/science/article/pii/S2667114X21000571

[20] R. Ben-Shachar, K. Koelle, Transmission-clearance trade-offs indicate that dengue virulence evolution depends on epidemiological context, Nature Communications 9. doi:10.1038/s41467-018-04595-w.

[21] N. J. White, Malaria parasite clearance, Malaria Journal 16. doi:10.1186/s12936-017-1731-1.

[22] J. M. Mansuy, C. Mengelle, C. Pasquier, S. Chapuy-Regaud, P. Delobel, G. Martin-Blondel, J. Izopet, Zika Virus Infection and Prolonged Viremiain Whole-Blood Specimens, Emerging Infectious Diseases 23 (5) (2017) 863–865. doi:10.3201/eid2305.161631. URL http://wwwnc.cdc.gov/eid/article/23/5/16-1631_article.htm

[23] C. Triplett, S. Dufek, N. Niemuth, D. Kobs, C. Cirimotich, K. Mack, D. Sanford, Onset and Progression of Infection Based on Viral Loads inRhesus Macaques Exposed to Zika Virus, Applied Microbiology 2 (3) (2022) 544–553. doi:10.3390/applmicrobiol2030042. URL https://www.mdpi.com/2673-8007/2/3/42

[24] Y. M. Tun, P. Charunwatthana, C. Duangdee, J. Satayarak, S. Suthisawat, O. Likhit, D. Lakhotia, N. Kosoltanapiwat, P. Sukphopetch, K. Boonnak, Virological, Serological and Clinical Analysis of Chikungunya Virus Infec-tion in Thai Patients, Viruses 14 (8) (2022) 1805. doi:10.3390/v14081805. URL https://www.mdpi.com/1999-4915/14/8/1805

